# *Sulfolobus acidocaldarius* adhesion pili power twitching motility in the absence of a dedicated retraction ATPase

**DOI:** 10.1101/2023.08.04.552066

**Authors:** Arthur Charles-Orszag, Marleen van Wolferen, Samuel J. Lord, Sonja-Verena Albers, R. Dyche Mullins

## Abstract

Type IV pili are ancient and widespread filamentous organelles found in most bacterial and archaeal phyla where they support a wide range of functions, including substrate adhesion, DNA uptake, self aggregation, and cell motility. In most bacteria, PilT-family ATPases disassemble adhesion pili, causing them to rapidly retract and produce twitching motility, important for surface colonization. As archaea do not possess homologs of PilT, it was thought that archaeal pili cannot retract. Here, we employ live-cell imaging under native conditions (75°C and pH 2), together with automated single-cell tracking, high-temperature fluorescence imaging, and genetic manipulation to demonstrate that *S. acidocaldarius* exhibits bona fide twitching motility, and that this behavior depends specifically on retractable adhesion pili. Our results demonstrate that archaeal adhesion pili are capable of retraction in the absence of a PilT retraction ATPase and suggests that the ancestral type IV pilus machinery in the last universal common ancestor (LUCA) relied on such a bifunctional ATPase for both extension and retraction.

## Introduction

Type IV pili are polymeric fibers that project from the outer surface of most archaea and bacteria (Pohlschroder *et al*., 2011; Berry and Pelicic, 2015; Chaudhury *et al*., 2018) where they function as an important interface between these cells and their environment. For example, type IV archaeal pili mediate adhesion to biotic and abiotic surfaces, inter- and intraspecies cell-cell interactions, biofilm formation, cell motility, and uptake of DNA in naturally competent species. Type IV pili are also receptors for bacteriophages and archaeal viruses (Bradley, 1974; Quemin *et al*., 2013; Hartman *et al*., 2019; Rowland *et al*., 2020; Wang *et al*., 2020) as well as central virulence factors in a number of plant and human bacterial pathogens. Despite decades of work, however, the type IV pili of archaea remain much less understood than their bacterial counterparts.

Similar to their bacterial counterparts, archaeal type IV pili are assembled from one or two major pilin proteins that polymerize into filaments. Prepilins contain a signature N-terminal sequence (the class III signal peptide) and are converted to mature pilins by the prepilin peptidase, PibD (Szabó *et al*., 2006; Szabó *et al*., 2007). Type IV pilus assembly machinery in archaea appear to comprise fewer components than those of bacteria. Typically, operons encoding the assembly factors contain genes for: (i) the assembly platform protein, (ii) an integral membrane protein, and (iii) the secretion/ assembly ATPase. Depending on the type of pilus formed, additional genes may be present that encode for proteins of unknown function. Phylogenetic analysis revealed that specific S-layer proteins are present close to these type IV pili clusters which might be important for the integration of the pilus into the archaeal cell envelope (Makarova *et al*., 2016).

In bacteria, the antagonist actions of two conserved ATPases, PilB and PilT, form the basis of type IV pilus dynamics. While PilB promotes fiber assembly, ATP hydrolysis by PilT fuels pilus disassembly and retraction. PilT-mediated retraction of a single pilus can generate forces of ∼100 pN and drive retraction at speeds of up to 1 μm per second (Merz *et al*., 2000; Maier *et al*., 2002). In bacteria, this dynamic behavior of type IV pili generates a characteristic mode of movement along the surface, called “twitching” motility, which is characterized by non-directional, saltatory motions of surface-adhered cells. Twitching motility is essential to surface colonization, biofilm formation, and dynamic self-aggregation (O’Toole and Kolter, 1998; Bonazzi *et al*., 2018; Gomez *et al*., 2023). For many years PilT was thought to be essential for retraction of type IV pili, leading to the proposal that archaeal pili are not capable of retraction and do not employ twitching motility (Chaudhury *et al*., 2018; Denise *et al*., 2019; Beeby, 2019). Recent work, however, has revealed the existence of PilT-independent retraction mechanisms in at least some bacteria (Ellison *et al*., 2017; Ellison *et al*., 2019; Hughes *et al*., 2022). While performing live-cell, time-lapse imaging of *S. acidocaldarius* at their normal growth temperature (75°C) we observed a type of surface-motility reminiscent of bacterial twitching motility (Charles-Orszag *et al*., 2021), challenging the idea that type IV pili in these cells are unable to retract.

The model hyperthermophilic crenarchaeon *Sulfolobus acidocaldarius* encodes three different type IV pili systems that are well studied. One is the archaellum, a motility structure, which is assembled via a type IV pili assembly mechanism, but which —in contrast to other type IV pili— rotates to generate torque for swimming motility (Jarrell and Albers, 2012; Tsai *et al*., 2020). The second are the UV-inducible pili (Ups), which initiate species-specific, cell-cell contacts that enable DNA exchange and DNA repair in Sulfolobales (Fröls *et al*., 2008; Fröls *et al*., 2008; van Wolferen *et al*., 2020). The third is the adhesive pilus called aap pilus (Henche *et al*., 2012a). This pilus is important for adhesion to surfaces and formation of biofilms (Henche *et al*., 2012b) and can be targeted by viruses as a cell adhesion factor (Quemin *et al*., 2013; Hartman *et al*., 2019; Rowland *et al*., 2020; Wang *et al*., 2020).

In this work, we characterize surface motility of wildtype *S. acidocaldarius* cells as well as several mutant strains lacking different cell surface appendages. Automated cell tracking in live-cell imaging assays at high temperature revealed that cells are capable of *bona fide* twitching motility, and that adhesion pili are specifically responsible for this phenomenon. Moreover, using super-resolution, live-cell, fluorescence imaging at high temperature we directly observed the rapid retraction of adhesion pili, and the coupling of these retraction events to saltatory cell movements. Finally, to determine the generality of these results, we analyzed cell motility in other Sulfolobales species and discovered that twitching motility is not unique to *S. acidocaldarius*. Our results indicate that PilT-independent pilus retraction might be common in archaea and strongly suggest that the ancestor of type IV pili in the last universal common ancestor (LUCA) already employed a single, bifunctional ATPase to drive both elongation and retraction and was capable of twitching motility.

## Results

### Adhesion pili mediate twitching motility in *S. acidocaldarius*

To study the motility of *S. acidocaldarius*, we transferred cells from exponentially growing cultures to an environmentally controlled microscope chamber heated to 75°C and performed timelapse imaging using Differential Interference Contrast (DIC) microscopy. In these experiments, we imaged microscope fields containing multiple cells for five minutes at one frame per second. We used automated segmentation and tracking with a trainable detector to identify and follow the movement of multiple cells in each timelapse movie. This machine learning approach enabled us to track several cells per field over hundreds of frames. The resulting tracks were then analyzed (Fig. 1).

**Figure 1.**
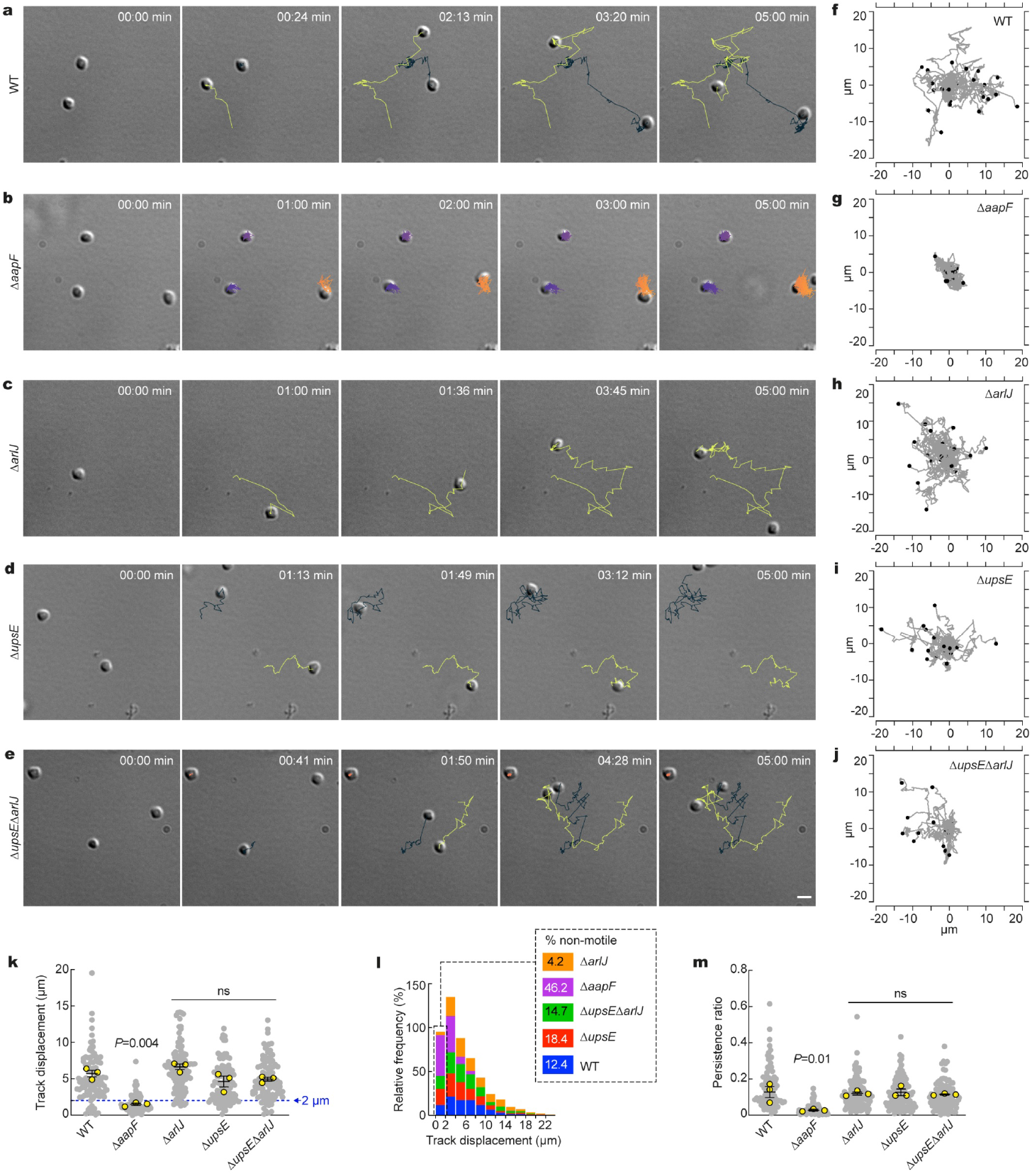
Automated tracking of type IV pili mutants in *S. acidocaldarius* at high temperature. **a-e**. Differential interference contrast (DIC) live-cell imaging of indicated *S. acidocaldarius* strains at 75°C. Shown are selected stills representing noticeable steps of glass-adhered cells over a five-minute observation window. Overlaid tracks were automatically obtained with TrackMate 7 after machine-trained detection of the cells with Weka. **f-j**. Worm plots of 15-30 representative tracks for each indicated strain. **k**. Track displacement measured over five minutes for each indicated strain. **l**. Distribution of individual track displacements in the indicated strains. Tracks with a displacement under 2 μm correspond to non-motile cells. **m**. Track persistence ratio in the indicated strains. Grey scatter dot plots correspond to the total number of cells analyzed (N = 200 WT, 222 Δ*aapF*, 284 Δ*arlJ*, 171 Δ*upsE*, 245 Δ*upsE*Δ*arlJ*). Superimposed yellow circles represent average values from each of n = 3 independent experiments. Bars represent the mean ± SEM of those three biological replicates.

As we previously observed for wild-type *S. acidocaldarius* (Charles-Orszag *et al*., 2021), glass-adhered cells from the parental strain (MW001, herein “WT”) were motile. Surface motility in WT cells appeared to be a saltatory random walk (Fig. 1a and f, and Supplementary Video 1). The average track displacement over 5 minutes was 5.7±0.44 μm. Not all cells were motile, however, with 12.4% of tracks having a total displacement less than 2 μm (the observed mean cell diameter was 1.6 μm under these experimental conditions (Charles-Orszag *et al*., 2021)) (Fig. 1k, l). WT tracks showed an average persistence ratio of 0.12±0.03, consistent with the apparent lack of directionality (Fig. 1m). For comparison, primary rat astrocytes exhibit a persistence ratio of 0.75 in a wound healing assay (De Pascalis *et al*., 2018). Therefore, the low persistence ratios measured here are consistent with a random walk rather than a directed, chemotaxis-like motility.

To determine which type IV pilus subtypes are involved in twitching motility, we analyzed surface motility in deletion strains missing either the archaellum membrane assembly protein ArlJ (Δ*arlJ*) or the UV-inducible pili assembly ATPase UpsE (Δ*upsE*). We also characterized the behavior of the double-deficient strain, Δ*upsE*Δ*arlJ*. All of these mutant strains exhibited motility indistinguishable from wildtype cells, with average total displacements of 4.6-6.6 μm (Fig. 1c-e, h-j, and k) and persistence ratios of 0.11-0.12 (Fig. 1m). Interestingly, while Δ*upsE* and Δ*upsE*Δ*arlJ* had a proportion of non-motile cells comparable to that of the WT (18.4% and 14.7% respectively) nearly all Δ*arlJ* cells were motile, with only 2% of tracks having a displacement below 2 μm (Fig. 1k, l). In sharp contrast, deleting the adhesion pilus assembly protein AapF (Δ*aapF*) appeared to abolish twitching motility, with surface-adhered cells merely exhibiting a continuous wobbly motion around a fixed point (Fig. 1b and g). Accordingly, track analysis showed that 46.2% of Δ*aapF* cells had a total displacement below 2 μm (Fig. 1k, l). Average track displacement (1.49 μm) and persistence ratio (0.02) in Δ*aapF* cells were both significantly smaller than in WT cells (Fig. 1k and m), confirming that Aap-deficient cells are non-motile.

Given the saltatory and non-directional nature of this mode of motility, as opposed to swimming, our data show that *S. acidocaldarius* exhibits *bona fide* twitching motility as seen in type IV pili-expressing bacteria. Taken together, these results demonstrate that twitching motility in *S. acidocaldarius* is specifically mediated by adhesion pili (Aap) and not by archaella or UV-inducible pili.

### Twitching motility in *S. acidocaldarius* is powered by retractile adhesion pili

Since a dedicated PilT-like retraction ATPase is absent in archaea, we sought to visualize type IV pili dynamics in live *S. acidocaldarius* cells to determine the mechanism by which Aap pili mediate surface motility. To do so, we non-specifically labeled surface-exposed lysines of membrane-associated proteins with a N-Hydroxysuccinimide (NHS)-ester functionalized Alexa Fluor dye, and imaged at high-temperature (Fig. 2). Labeled cells adhered to the glass surface and underwent twitching motility, indicating that Aap pili were functional. In addition to a bright fluorescent signal originating from cell wall proteins, we detected Aap pili as thin, surface-attached fibers of up to 11 μm long. Most importantly, cell movement coincided with rapid retraction of Aap pili in two ways: cells either moved forward in the direction of pilus retraction or moved backward upon loss of surface attachment after a pilus retracted (Fig. 2b, d and Supplementary Video 2). The retraction of Aap pili also supported dynamic direct cell-cell interactions (Fig. 2b). In cells where adhesion was mediated by a single adhesion pilus, the average pilus retraction speed was 2 μm.sec^-1^. The speed of retraction did not correlate to pilus length (Fig. 2c).

**Figure 2.**
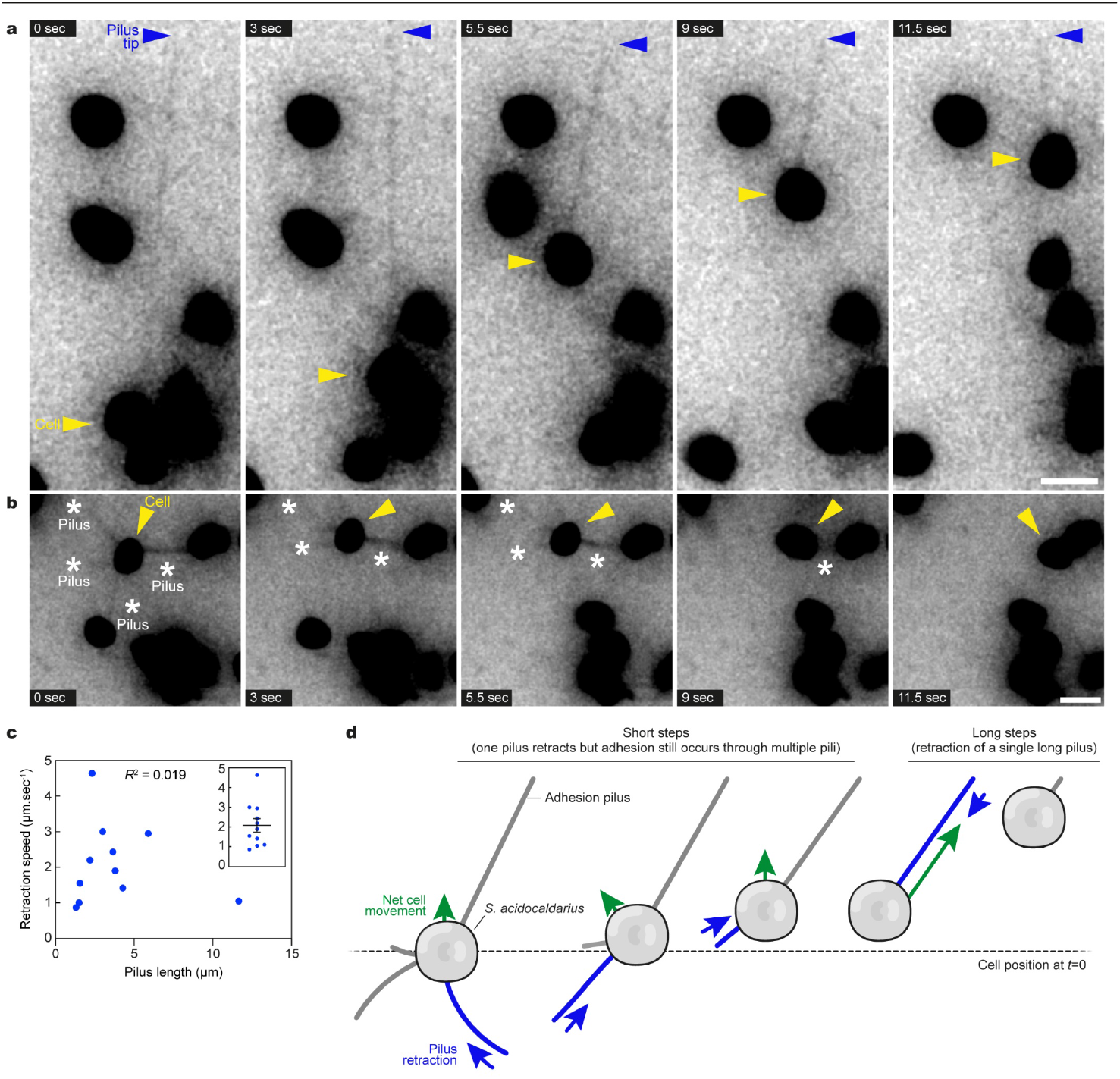
Super-resolution fluorescence imaging of *S. acidocaldarius* adhesion pili dynamics at high temperature. Surface proteins of wild-type *S. acidocaldarius* cells were labeled non-specifically with AlexaFluor 568 NHS-ester. Glass-adhered cells were imaged in Structured Illumination Microscopy (iSIM) every 500 ms. Shown are selected stills from representative movies. **a**. A cell (yellow arrowhead) with a pilus of 11 μm (pilus tip denoted by blue arrowhead) moves across the surface as the pilus gets shorter. **b**. A cell (yellow arrowhead) with four visible pili (white asterisks) connected to two neighbor cells. The cell moves in short steps as the first pili retract, and finally gets in close contact with one of the neighboring cells upon retraction of the last pilus. Scale bars, 2 μm. **c**. Pilus retraction speed was measured in 11 cells in which adhesion was mediated by a single adhesion pilus, and plotted against pilus length. The inset represents the mean retraction speed ± SEM. **d**. Interpretative diagram showing how retraction of adhesion pili mediates short and long steps observed in the twitching motility of *S. acidocaldarius*.

These results demonstrate that *S. acidocaldarius* Aap pili are capable of retraction, and that pilus retraction powers twitching motility in these cells.

### Role of the minor pilin AapA

To investigate the molecular mechanisms of Aap pilus retraction in more detail, we investigated twitching in *S. acidocaldarius* strains deficient for other genes in the *aap* operon (Fig. 3a). In addition to the major pilin AapB, the ATPase AapE, and the membrane protein AapF, the *aap* pilus operon comprises two other genes, *aapA* and *aapX*, whose functions are unclear (Fig. 3a). AapX is annotated as an iron-sulfur oxidoreductase, and deletion of the *aapX* gene leads to a lack of pilus assembly (Henche *et al*., 2012a). AapA is a type IV pilin that contains a class III signal peptide that is processed by the prepilin peptidase PibD, and it was proposed to be a minor pilin that might co-assemble in the pilus fiber with AapB. AapA was first thought to be indispensable for pilus assembly (Henche *et al*., 2012a), but a recent study demonstrated that a Δ*aapA* mutant still assembles pili (Gaines *et al. &* Wang *et al*., bioRxiv 2023), suggesting that AapA might not co-polymerize with AapB in the pilus fiber, and that AapA does not have a function in pilus assembly.

**Figure 3.**
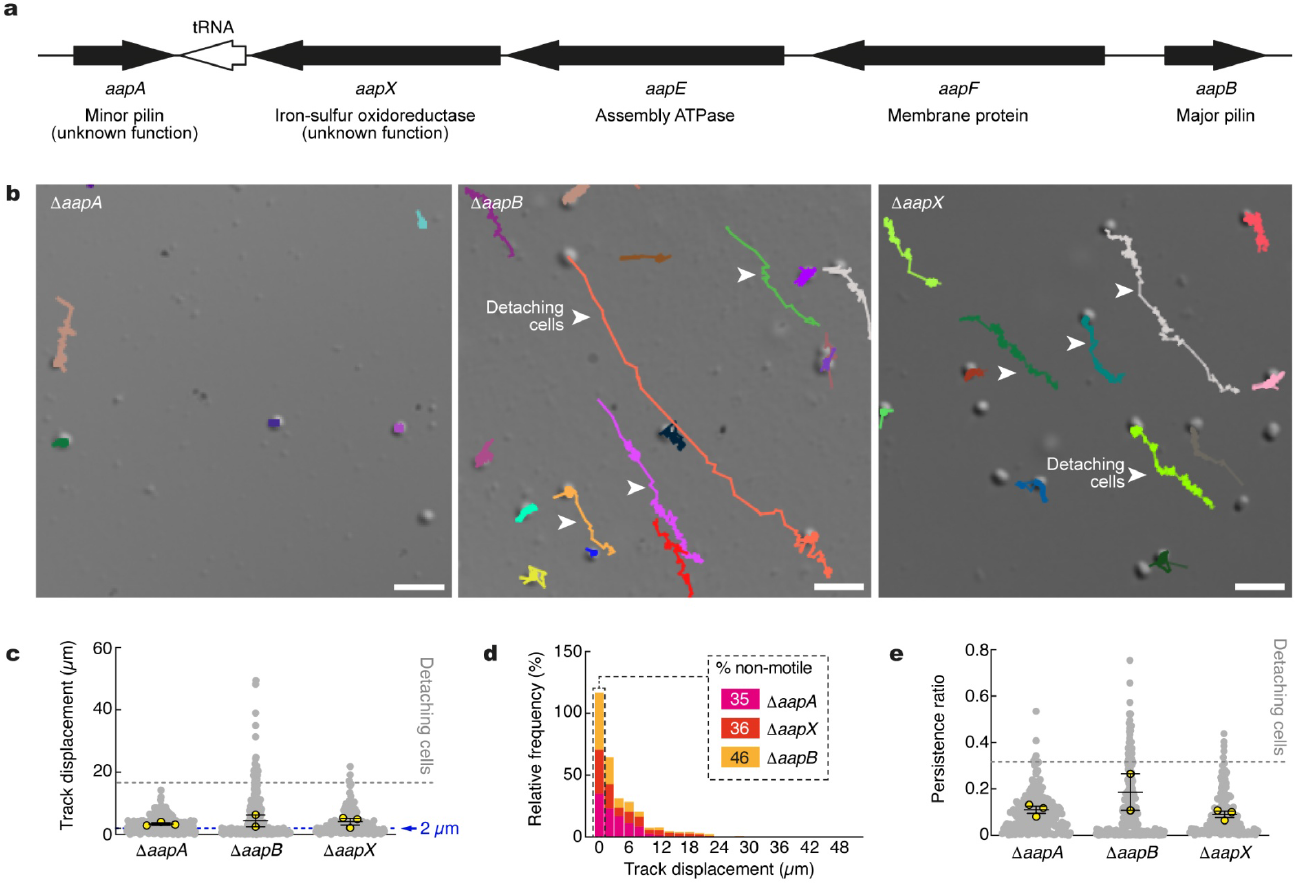
Automated tracking of *S. acidocaldarius* Aap mutants. **a**. Map of the Aap pilus operon in *S. acidocaldarius*. **b**. Representative examples of tracks generated for indicated Aap mutants. Arrowheads denote long aligned tracks generated by detaching cells. Scale bars = 5 μm. **c**.Track displacement measured over five minutes for each indicated strain (Δ*aapA*: N = 418, Δ*aapB*: N = 302, Δ*aapX*: N = 427). **d**. Distribution of individual track displacements in the indicated strains. Tracks with a displacement <2 μm were defined as non-motile. **e**. Track persistence ratio in the indicated strains. Superimposed yellow circles represent sample means from each of n=3 (Δ*aapA*, Δ*aapX*) or n=2 (Δ*aapB*) independent experiments. Bars represent the mean ± SEM of those biological replicates.

Surprisingly, the Δ*aapA*, Δ*aapB*, or Δ*aapX* deletion mutants all exhibited drastically reduced twitching motility in our live-cell imaging assay (Fig. 3). Similar to the Δ*aapF* mutant cells (Fig. 1), most cells were non-motile (Fig. 3b, c and d). However, in Δ*aapB*, and to a lesser extent in Δ*aapX*, some cells appeared very loosely attached to the surface. As a result, these cells would detach and roll away over long distances because of the high convection flows in the chamber (Fig. 3b), generating long tracks (Fig. 3c) with a higher persistence ratio (Fig. 3e).

These data are consistent with previous literature showing that Δ*aapB* and Δ*aapX* do not assemble pili. However, although the putative minor pilin AapA appears to not be involved in pilus formation, our results suggest that AapA plays an important role in pilus retraction and twitching motility.

### Twitching motility is present in other Sulfolobales

Other members of the Sulfolobales encode an Aap pilus operon (Henche *et al*., 2012a). To test whether type IV pilus-mediated twitching motility is specific to *S. acidocaldarius*, we investigated cell motility in other Sulfolobale species that express adhesion pili. Automated tracking of cells imaged at high temperature showed that *Sa. solfataricus* and *S. islandicus* M16 and REY15A all include some cells that exhibited twitching motility, although with a lower average track displacement and with a higher proportion of non-motile cells overall compared to *S. acidocaldarius* (Fig. 4 and Supplementary Video 4).

**Figure 4.**
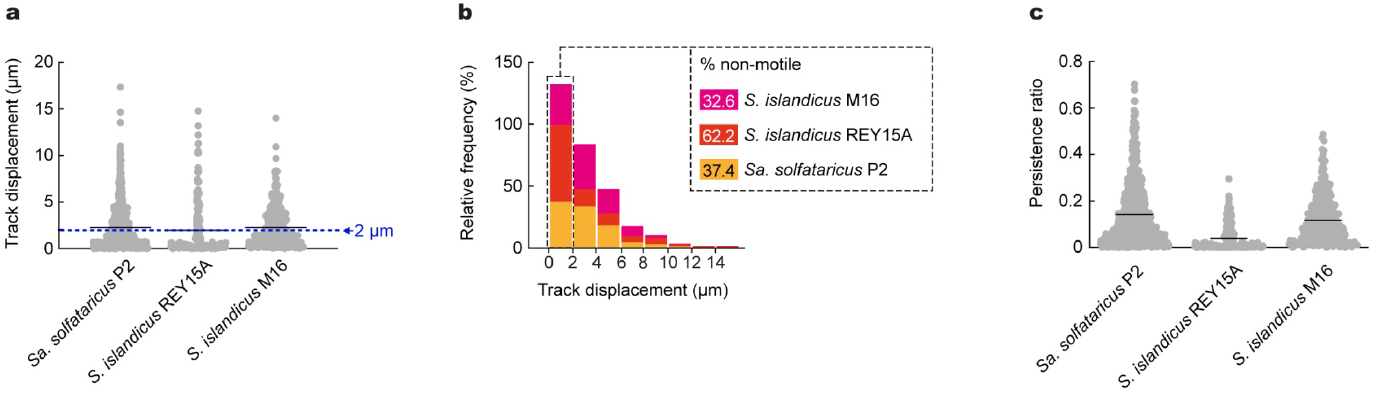
Automated tracking in other Sulfolobales species. **a**.Track displacement and persistence ratio measured over five minutes for each indicated species (*Saccharolobus solfataricus* P2: N = 836, *Sulfolobus islandicus* REY15A: N = 180, *Sulfolobus islandicus* M16: N = 344. **b**. Distribution of individual track displacements in the indicated species. Tracks with a displacement under 2 μm correspond to the percentage of non-motile cells. **c**. Track persistence ratio in the indicated species.

Our data strongly suggests that twitching motility is also supported by type IV pili in *Sa. saccharolobus* and *S. islandicus*, and demonstrates that twitching motility is not restricted to *S. acidocaldarius* but common in Sulfolobales. This, in turn, indicates that PilT-independent type IV pilus retraction is widespread among archaea.

## Discussion

At any given moment most prokaryotes on planet Earth are forming part of a biofilm (Flemming *et al*., 2016; van Wolferen *et al*., 2018) often comprising multiple species that either coexist in synergy or compete for resources, making these environments key drivers of evolution. The formation of biofilms is dependent on the ability of cells to adhere to, explore, and eventually colonize surfaces. In both bacteria and archaea, type IV pili are central players of biofilm formation as they promote surface attachment. In most bacteria, the antagonist actions of the conserved ATPases PilB and PilT promote assembly and retraction of pilus fibers, thereby powering twitching motility which is essential for surface exploration and dynamic cell-cell interactions. The type IV pilus assembly machinery is ubiquitous in prokaryotes and was likely present in the last universal common ancestor, LUCA (Berry and Pelicic, 2015; Makarova *et al*., 2016; Denise *et al*., 2019). However, the retraction ATPase, PilT, is only found in bacteria, leading to the assumption that archaea are not capable of twitching motility (Chaudhury *et al*., 2018; Denise *et al*., 2019; Beeby, 2019). However, recent observations of live *S. acidocaldarius* cells challenged this concept as the cells appeared highly surface-motile, in a manner reminiscent of bacterial twitching motility (Charles-Orszag *et al*., 2021).

In the present work, we show that *S. acidocaldarius* cells deficient for adhesion pili are no longer surface-motile, whereas deletion of any of the other type IV pili subtypes (archaella and UV-inducible pili) did not impair surface motility. Given the saltatory and non-directional mode of motility in WT cells, these results demonstrate that *S. acidocaldarius* uses twitching motility to adhere to and explore surfaces, and that this behavior is specifically mediated by adhesion pili. Of note, our data shows that cells deficient for archaella assembly (Δ*arlJ*) have a higher proportion of motile cells compared to the WT, suggesting the existence of a genetic crosstalk between type IV pilus subtypes. Indeed, it was previously observed that adhesion pili are upregulated in a Δ*arlJ* mutant (Henche *et al*., 2012b).

The fact that *S. acidocaldarius* adhesion pili allow twitching motility implies that adhesion pili are dynamic, either via retraction or via shedding. However, as described earlier, archaea do not express homologs of bacterial PilT ATPases responsible for pilus retraction. Here, super-resolution fluorescence microscopy of living cells clearly shows that cells move around by retracting surface-adhered pili. There, pili appear as thin extended threads that seem adhered through the pilus tip, as proposed for some bacteria (Kennouche *et al*., 2019). Upon pilus retraction, *Sulfolobus* cells can move in the direction of retraction. In the case where cells are attached to the surface through multiple pili, they move in a direction that brings the cell body in the middle of the remaining adhered pili. These results demonstrate that adhesion pili are retractile.

Two bacterial species, *Caulobacter crescentus* and *Vibrio cholerae*, are also known to lack a retraction ATPase and it was shown that the assembly ATPase PilB is responsible for both pilus assembly and retraction (Ellison *et al*., 2017; Ellison *et al*., 2019; Hughes *et al*., 2022). In our case, it is likely that the adhesion pili assembly ATPase AapE also coordinates pilus retraction. Of note, the speed of pilus retraction in *S. acidocaldarius* is comparable to that of bacterial pili, i.e. ∼1 μm.sec^-1^ (Maier *et al*., 2002). It was difficult to prove that AapE is involved in the retraction as deletion of AapE leads to a lack of assembly of Aap pili (Henche *et al*., 2012a). We attempted to overexpress AapE Walker A and B mutants from plasmids in wild-type cells to observe whether Aap pili could still retract in the conditions, but these experiments proved difficult to interpret since very tight inducible promoters do not exist for expression of proteins in *Sulfolobus* species.

In addition, our data reveal that cells deficient for the minor pilin AapA are incapable of twitching. However, in a new study that used cryo-electron microscopy to determine the structure of the Aap pilus, the authors found that the pilus fiber is exclusively composed of subunits of the major pilin AapB (Gaines *et al. &* Wang *et al*., bioRxiv 2023). Even in bacteria, the function of minor pilins has been subject to debate, and studies in some bacterial species have shown that minor pilins actually exert their function from the cytoplasmic membrane and not from the pilus fiber (Imhaus and Duménil, 2014). Our results suggest that the minor pilin AapA plays a role in twitching motility. One possibility would be that AapA is present at the pilus tip in minute amounts and mediates substrate adhesion and transduction of mechanical signals. Alternatively, AapA could function in the cytoplasmic membrane and mediate the transition from the assembly to the retraction activity of the AapE ATPase. Therefore, more work is required to decipher the molecular mechanism by which AapE switches between pilus assembly and retraction in *Sulfolobus*.

Interestingly, it was proposed that the retraction ATPase PilT evolved in bacteria as a result of a gene duplication of the assembly ATPase PilB, followed by a separation of the polymerization and retraction activities between the two ATPases (Denise *et al*., 2019; Beeby, 2019). In this work, we show that multiple Sulfolobales species are capable of twitching motility, suggesting that PilT-independent pilus retraction might be a common mechanism in archaea. However, it is worth noting that different species might present different abilities to twitch. For example, twitching motility was not been observed in the piliated halophile *Haloferax volcanii* (Odermatt *et al*., 2023). In the present study, we show that *Sa. solfataricus* and *S. islandicus* twitch less efficiently than *S. acidocaldarius*. Perhaps this reflects adaptation to specific ecological niches. For instance, as opposed to *Sa. solfataricus* and *S. islandicus*, it is thought that *S. acidocaldarius* preferentially colonizes hot springs rocky periphery, possibly thanks to a more robust and/or active Aap pilus system (Cadillo-Quiroz *et al*., 2012; Henche *et al*., 2012a), which is in good accordance with the ability of *S. acidocaldarius* to form thicker and more complex biofilm structures compared to other Sulfolobales (Henche *et al*., 2012b).

Along with driving cell motility, pilus retraction in bacterial species such as *E. coli, C. crescentus* and *P. aeruginosa* can be hijacked by bacteriophages, which use it to move into close proximity with the host cell surface (Bradley, 1972a; Bradley, 1972b; Jacobson, 1972; Bradley, 1974; Sommer and Newton, 1988; Skerker, 2000). In archaea too, type IV pili can serve as receptors for archaeal viruses (Quemin *et al*., 2013; Hartman *et al*., 2019; Rowland *et al*., 2020; Wang *et al*., 2020). Our study, therefore, suggests that archaeal viruses might utilize type IV pilus retraction to infect their host, in a way similar to bacteriophages.

**Table 1.** List of strains used in this study.

| Strain | Description | Source |
| --- | --- | --- |
| MW001 | ΔpyrEF (91–412 bp); parental strain | Wagner *et al.*, 2012 |
| MW109 | ΔupsE (deletion of *saci1494* in MW001) | Wagner *et al.*, 2012 |
| MW153 | ΔaapA (deletion of *saci1174* in MW001) | Henche *et al.*, 2012a |
| MW154 | ΔaapB (deletion of *saci1179* in MW001) | Henche *et al.*, 2012a |
| MW155 | ΔarlJ (formerly *flaJ*, deletion of *saci1172* in MW001) | Henche *et al.*, 2012b |
| MW156 | ΔaapF (deletion of *saci2318* in MW001) | Henche *et al.*, 2012b |
| MW158 | ΔarlJ ΔupsE | Henche *et al.*, 2012b |
| MW159 | ΔaapX (deletion of *saci2316* in MW001) | Henche *et al.*, 2012a |
| MW160 | ΔaapE (deletion of *saci2317* in MW001) | Henche *et al.*, 2012a |
| *Saccharolobus solfataricus* P2 |  | DSMZ |
| *Sulfolobus islandicus* REY15A |  | Gift from Quinxin She |
| *Sulfolobus islandicus* M16 |  | Gift from Rachel Whitaker |

In summary, our work suggests that the ancestor of the type IV pius machinery in LUCA relied on a single bifunctional ATPase for both pilus assembly and retraction. This in turn implies that LUCA was already able to dynamically explore and colonize surfaces the same way modern prokaryotes do, underlying the evolutionary success of biofilms as a prokaryotic lifestyle.

## Materials and methods

### Cell culture

*Sulfolobus acidocaldarius, Sulfolobus islandicus* and *Saccharolobus solfataricus* were grown aerobically at 75°C with shaking in Brock’s minimal media (Brock *et al*., 1972). *S. acidocaldarius* was grown in the presence of 20 μg.mL^-1^ uracil (Sigma-Aldrich).

### Automated cell tracking

*S. acidocaldarius* cells from exponentially growing cultures were imaged in a DeltaT cell micro-environment control system (Bioptechs) at 75°C as previously described (Charles-Orszag *et al*., 2021) with the difference that the glass coverslip was not modified with solidified media so cells were allowed to adhere to glass, or in a VAHeat chamber controlled by the VAHeat system (Interherence) on a Nikon Ti-E inverted microscope equipped with a 100x 1.45NA PlanApo objective and a Point Grey CMOS camera (CM3-U3-50S5M-CS FLIR; 85×71 μm fields of view), or a Zeiss Observer Z1 microscope equipped with a Plan-Apochromat 100x 1.40NA objective (166×130 μm fields of view). Movies were recorded at a rate of 1 frame per second for a total of 5 min.

The first four frames from a representative movie were used to train the Weka detector (Witten *et al*., 2011) in Fiji (Schindelin *et al*., 2012) after decreasing the frame size so that the longest dimension was smaller than 1000 pixels. Weka was trained to detect surface-adhered cells (in focus) and ignore unadhered cells (out of focus) as well as signal from the background, or debris in the culture media. These categories were uploaded in TrakMate 7 (Ershov *et al*., 2022) operated in Fiji and cells were automatically tracked in full-length movies. Low-quality tracks, tracks corresponding to multiple cells wrongfully detected as one, or tracks including cell-cell interactions events were discarded by adjusting the detection threshold in TrackMate 7 and manual pruning. The dataset was refined by visual inspection and manual bridging of gaps within tracks that were interrupted by transient loss of focus or detection defects.

### Live cell fluorescence imaging of adhesion pili dynamics

10 mL of *S. acidocaldarius* cells from an exponentially growing culture were collected by centrifugation at 5000 g at room temperature. The pellet was washed once in phosphate buffered saline (PBS), resuspended in 225 μL PBS, and 25 μL of Alexa Fluor 568-NHS-ester dye (Thermo Scientific) freshly reconstituted at 0.5 μg.mL^-1^ in anhydrous DMSO was added. Cells were incubated with the dye for 15 min protected from light, with gentle shaking, at room temperature. Cells were then pelleted, washed twice with minimal Brock’s media, and recovered in 5 mL complete Brock’s media at 75°C with shaking for 1.5 h prior to imaging.

Cells were imaged in a VAHeat chamber controlled by the VAHeat system (Interherence) on a Nikon Ti-E inverted microscope equipped with a 100x 1.45NA PlanApo objective and an additional 1.5X intermediate tube lens, 561 nm laser, Hamamatsu Quest camera, and a VT-iSIM super-resolution module. Movies were recorded at a rate of two frames per second.

## Data analysis

Track analysis and image processing for figure preparation were made in Fiji (Schindelin *et al*., 2012). Data analysis and preparation of SuperPlots (Lord *et al*., 2020) were done in Prism (GraphPad). Figures and illustrations were prepared in Adobe Illustrator.

## Acknowledgments

This work was supported by the National Institute of General Medical Sciences of the National Institutes of Health (R01-GM061010 to RDM), by the Howard Hughes Medical Institute Investigator Program (RDM), and by a Momentum grant (VW Foundation grant number 94933 to SVA and MVW). ACO is a Simons Fellow of the Life Sciences Research Foundation.

## Author contributions

ACO: conceptualization, methodology, investigation, formal analysis, validation, visualization, figure preparation, and writing original draft preparation, and review and editing. MVW: conceptualization, methodology, investigation, formal analysis, validation, visualization, and review and editing. SL: methodology, formal analysis, validation, visualization, figure preparation, writing original draft preparation, and review and editing. SVA & RDM: funding acquisition, supervision, methodology, investigation, formal analysis, writing original draft preparation, and review and editing. All authors contributed to the article and approved the submitted version.

## Competing interests

The authors declare that the research was conducted in the absence of any commercial or financial relationships that could be construed as a potential conflict of interest.

## Supplementary Information

**Supplementary Figure 1.**
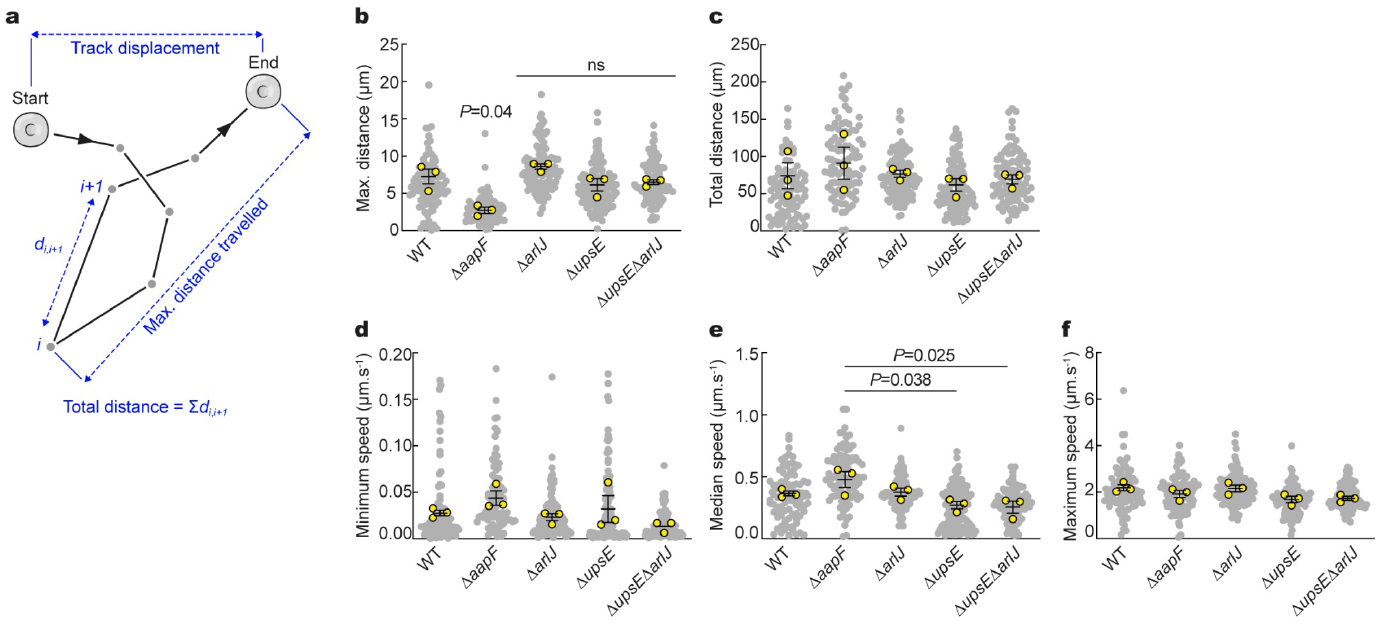
Automated tracking of type IV pili mutants in *S. acidocaldarius* at high temperature (continued). **a**. Diagram representing a typical track obtained from a cell trajectory in TrackMate 7, and illustrating the biological parameters that were measured in the present study. Adapted from https://imagej.net/plugins/trackmate/algorithms. **b**. Maximum distance travelled over five minute for each indicated strain. **c**. Total distance travelled over five minutes for each indicated strain. **d-f**. Minimum, median and maximum track speeds in each indicated strain. Grey scatter dot plots correspond to the total number of cells analyzed (200 WT, 222 Δ*aapF*, 284 Δ*arlJ*, 171 Δ*upsE*, 245 Δ*upsE*Δ*arlJ*). Superimposed yellow circles represent average values from each of n=3 independent experiments. Bars represent the mean ± SEM of those three biological replicates.

**Supplementary Video 1**

Automated tracking of type IV pili mutants in *S. acidocaldarius* at high temperature. Differential interference contrast (DIC) live-cell imaging of indicated *S. acidocaldarius* strains at 75°C. Tracks of each cell are overlaid in a different color.

**Supplementary Video 2**

Super-resolution fluorescence imaging of *S. acidocaldarius* adhesion pili dynamics at high temperature. Surface proteins of wild-type *S. acidocaldarius* cells were labeled non-specifically with Alexa Fluor 568 NHS-ester. Glass-adhered cells were imaged in instant Structured Illumination Microscopy (iSIM) every 500 ms.

**Supplementary Video 3**

Automated tracking of *S. acidocaldarius* Aap mutants. Differential interference contrast (DIC) live-cell imaging of indicated species at 75°C. Tracks of each cell are overlaid in a different color.

**Supplementary Video 4**

Automated tracking of different Sulfolobales species at high temperature. Differential interference contrast (DIC) live-cell imaging of indicated species at 75°C. Tracks of each cell are overlaid in a different color.

## Notes

### Competing Interest Statement

The authors have declared no competing interest.

